# Dynamic and Biphasic Regulation of Cell Migration by Ras

**DOI:** 10.1101/2025.02.13.638204

**Authors:** Yiyan Lin, Eleana Parajón, Qinling Yuan, Siyu Ye, Guanghui Qin, Yu Deng, Jane Borleis, Ariel Koyfman, Pablo A. Iglesias, Konstantinos Konstantopoulos, Douglas N. Robinson, Peter N. Devreotes

## Abstract

Ras has traditionally been regarded as a positive regulator and therapeutic target due to its role in cell proliferation, but recent findings indicate a more nuanced role in cell migration, where suppressed Ras activity can unexpectedly promote migration. To clarify this complexity, we systematically modulate Ras activity using various RasGEF and RasGAP proteins and assess their effects on migration dynamics. Leveraging optogenetics, we assess the immediate, non-transcriptional effects of Ras signaling on migration. Local RasGEF recruitment to the plasma membrane induces protrusions and new fronts to effectively guide migration, even in the absence of GPCR/G-protein signaling whereas global recruitment causes immediate cell spreading halting cell migration. Local RasGAP recruitment suppresses protrusions, generates new backs, and repels cells whereas global relocation either eliminates all protrusions to inhibit migration or preserves a single protrusion to maintain polarity. Consistent local and global increases or decreases in signal transduction and cytoskeletal activities accompany these morphological changes. Additionally, we performed cortical tension measurements and found that RasGEFs generally increase cortical tension while RasGAPs decrease it. Our results reveal a biphasic relationship between Ras activity and cellular dynamics, reinforcing our previous findings that optimal Ras activity and cortical tension are critical for efficient migration.

**Significance:** This study challenges the traditional view of Ras as solely a positive regulator of cell functions by controlling of gene expression. Using optogenetics to rapidly modulate Ras activity in *Dictyostelium*, we demonstrate a biphasic relationship between Ras activity and migration: both excessive and insufficient Ras activity impair cell movement. Importantly, these effects occur rapidly, independent of transcriptional changes, revealing the mechanism by which Ras controls cell migration. The findings suggest that optimal Ras activity and cortical tension are crucial for efficient migration, and that targeting Ras in cancer therapy should consider the cell’s initial state, aiming to push Ras activity outside the optimal range for migration. This nuanced understanding of the role of Ras in migration has significant implications for developing more effective cancer treatments, as simply inhibiting Ras might inadvertently promote metastasis in certain contexts.

## Introduction

Ras small GTPases are highly versatile proteins that play pivotal roles in numerous signaling events, regulating fundamental cell functions such as growth, survival, differentiation, and migration (1–3). A delicate balance of Ras activity is crucial for proper gene expression, cellular functions, and tissue development and dysregulation of Ras can lead to developmental abnormalities and the onset of diseases such as cancer (4–6). It is generally believed that Ras serves as a *positive* regulator of these functions, and Ras has garnered enormous attention as a therapeutic target, with most research focused on proliferation and differentiation (7–9). However, recent studies of cell migration have shown that effects of modulating Ras activity depend largely on the initial state of the cell. Studies in rapidly moving cells such as *Dictyostelium discoideum* and human neutrophils have shown that constitutively active Ras isoforms induce more protrusion formation as might be expected but excess protrusions can lead to cell spreading that precludes movement while dominant negative Ras isoforms or expression of RasGAPs that limit protrusions can promote cell polarization and migration (10, 11). These findings highlight the complexity of Ras signaling in cell migration and show that the relationship between Ras activity and cell migration is not straightforward. The counterintuitive outcomes must be considered when assessing Ras as a target to treat cancer. An inhibitor that limits growth may in fact promote metastasis. Therefore, it is important to obtain a full picture of how modulating Ras activity affects cell migration, as well as other aspects, such as cell morphology, macropinocytosis, and mechanical properties.

During growth factor stimulation, transcriptional and translational events are not immediately observable whereas the triggering of Ras/Raf/MEK/ERK and Ras/PI3K/mTOR signal transductions cascades are readily apparent. Cells undergo immediate morphological changes caused by dramatic cytoskeleton remodeling, leading to cell spreading, ruffles, and protrusion formation (12–15). Furthermore, optogenetically activating or inhibiting Ras can rapidly create new protrusions and guide cell migration or eliminate protrusions and halt migration, respectively, on timescales much faster than those required for changes in gene expression (10, 11).

Mutations in the human NF1 gene, a RasGAP, cause the severe genetic disorder neurofibromatosis type 1 (16, 17). The defects are typically associated with increased proliferation with little consideration of effects on cell morphology and dynamics (18, 19). However, macropinocytosis is upregulated in Ras-driven tumors and the process appears to be essential for driving accelerated growth. Interestingly, activation of Ras isoforms is dynamically associated with macropinocytotic cups in axenic strains of the social amoeba *Dictyostelium* (20). This upregulation of macropinocytosis in these strains is caused by inactivation of a single NF1 homologue which leads to hyperactivated Ras activity and underlies the massive increase in macropinocytosis.

Surprisingly, cells with activated Ras do not grow dramatically faster, suggesting that G1 checkpoint control is not critical in these free-living amoebae. For these reasons, this model organism offers the possibility of studying the effects of Ras activity modulation on cell dynamics without complication from effects on proliferation.

Ras activity is regulated by guanine nucleotide exchange factors (RasGEFs) and GTPase-activating proteins (RasGAPs) (21, 22). The *Dictyostelium* genome includes about 25 genes encoding RasGEFs and 14 for RasGAPs (23). Among them, Aimless and GefR can activate RasC and RasG, and their loss-of-function mutants have defects in chemotaxis(24). GefB, GefM, GefU, and GefX are all reportedly important for random migration and spontaneous signaling events (23). RasGAPs C2GAP1 and C2GAP2 interact with RasC and RasG and are essential for GPCR-mediated Ras signaling in chemotaxis (25–27). NF1, NfaA and IqgC are RasGAPs that are mainly involved in the regulation of macropinocytosis or phagocytosis (20, 28–30). Previously, we found that overexpression or optogenetically recruitment of C2GAPB in vegetative *Dictyostelium* unexpectedly polarized cells, albeit its inhibitory role on Ras activity (10). Considering the complexity and importance of Ras signaling on cell shape and migration, we aim to systematically modulate Ras activity over a broad spectrum and further extend our investigation to GefA (Aimless) and NGAP (C2GAP1). Through overexpression or optogenetic manipulation, we sought to determine their effects on cell migration, cell morphology, signal transduction, macropinocytosis, as well as the mechanical properties of cells. Our evidence suggests a biphasic relationship between Ras activity and cell migration. By dissecting this critical node in the signaling network, we provide new insights into the coordinated regulation of cellular processes by Ras signaling pathways.

## RESULTS

### Exchange Factor GefA Alters Cell Morphology and Migration

Because Ras is thought to function as a positive regulator in cell migration and chemotaxis, we first examined the immediate effects of increased Ras activity on cellular protrusions and migration directions. To accomplish this aim, we designed an optogenetic system to recruit SspB fused GefA protein from the cytosol to the cell membrane anchored iLiD when blue light is presented(31–33). We hypothesized that recruiting more GefA protein at the front of cells would enlarge the protrusions, while recruiting it to the back would create new protrusions and reverse migration directions (**Fig. 1A**). When we locally illuminated a protrusion, the recruitment led to an expansion of the protrusion within 30 seconds and induced cells to shift the migration direction (**Fig. 1B**). Next, we attempted to make the cell reverse direction by recruiting GefA to the back. This manipulation did induce protrusions, but because vegetative *Dictyostelium* cells are less polarized, the direction changes were inconclusive. Therefore, we loaded cells into microchannel devices which enforce a defined direction. As shown in **Fig. 1C**, when GefA-expressing cells moved in one direction, we locally concentrated GefA at the trailing edge. In less than 3 minutes, the cell switched direction. We were able to repeatedly change directions by alternately recruiting GefA at different positions. The movement of the edge of the cell lagged the recruitment of the GefA by 20-60 sec (**Movie S1**). Compared to no light treatment where cells move to either left or right, the recruitment of GefA consistently guided cells move toward the side of GefA accumulation (**Fig. 1D**). Strikingly, local recruitment of GefA can induce new protrusions formation and guide the cell migration in Gβ-null cells, which are defective in most receptor-mediated signal transduction events and do not carry out chemotaxis (**Fig. S1A**).

**Fig. 1.**
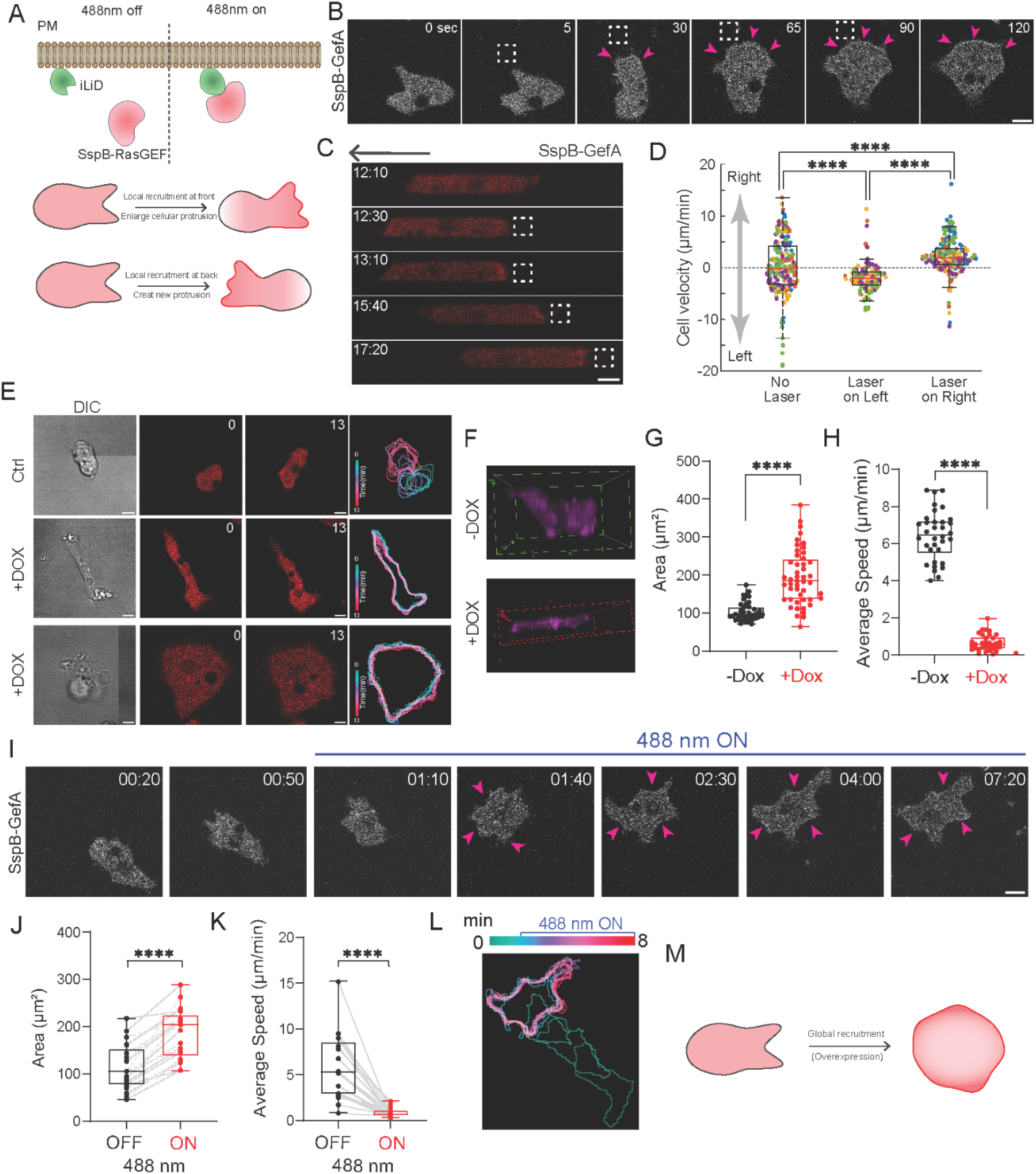
Activation of Ras by GefA alters cell morphology and migration. **(A)**Schematic of the optogenetics tool. Recruitment of Opto-GefA (mRFPmars-SspB-GefA) to membrane-anchored iLiD upon 488 nm illumination. Typical effects at cell front or back are indicated. **(B)** Opto-GefA recruited near a protrusion from migrating vegetative *Dictyostelium* cells on 2D glass surface with 488 nm light (dashed white box). Pink arrows highlight the expansion of the protrusion. Scale bars, 5 µm. **(C)** Opto-GefA recruited to the back of cell migrating to the left in a microchannel (dashed white box). Arrow indicated the migration direction before light stimulus. Scale bars, 5 µm. **(D)** Instantaneous velocity of Opto-GefA cells when the laser light is off, to the left or right of cells. Colored dots represent each of n=5 cells; total number of measurements are 215, 121, and 184, respectively. Asterisks indicate significant difference, ****P ≤ 0.0001 (two-sided Mann–Whitney test). **(E)** Time-lapse confocal images of vegetative *Dictyostelium* cells expressing empty vector control (red; top panel) or mRFPmars-GefA (red; middle and bottom panel) after overnight doxycycline (Dox) treatment. Time in minutes. Scale bars, 5 µm. DIC images (left-most panel) and overlaid outlines (color-coded 1-min intervals) of control or GefA-expressing cells (right-most panel). **(F)** 3D z-stack images of GefA-expressing cell minus (top) or plus (bottom) Dox induction. Box-and-whisker plots of cell area **(G)** and average cell speed **(H)** before (black; −DOX) or after (red; +DOX) overnight induction of GefA. n = 40 (−Dox) or n = 49 (+Dox) cells examined over three independent experiments; asterisks indicate significant difference, ****P ≤ 0.0001 (two-sided Mann–Whitney test). The boxes extend from 25th to 75th percentiles, median is at the center and whiskers and outliers are graphed according to Tukey’s convention (GraphPad Prism 8). **(I)** Snapshots of vegetative *Dictyostelium* cells expressing Opto-GefA on 2D glass surface. Opto-GefA is recruited globally by switching on 488 nm laser. Pink arrows highlight the expansion of the entire cell. Time in min:s format. Scale bars, 5 µm. Box-and-whisker plots of cell area **(J)** and average cell speed **(K)** before (black; 488 nm OFF) or after (red; 488 nm ON) global recruitment of GefA. n = 21 cells examined over three independent experiments. Statistical analysis as in **(G)** and **(H)**. **(L)** Color-coded (1-min interval) outlines of a representative cell before and after the presence of 488 nm light. **(M)** Cartoon summarizes the effects of global opto-GefA recruitment or overexpression of GefA.

Given the obvious effects of increasing Ras activity on promoting cellular protrusions and guiding cell migration direction, we hypothesized that overexpressing GefA would promote cell migration. Suspecting that inhibitory effects of continuous expression of active Ras proteins on cytokinesis selects for low expressors, we used an inducible system to control GefA expression. Axenic vegetative cells exhibit an amoeboid shape and often form several protrusions simultaneously. They are less polarized then developed cells, but nevertheless display moderate movement. However, upon the overnight induction of GefA expression, the cells exhibited either a large, flat, pancake-like single global protrusion or thin, elongated morphologies with multiple opposing protrusions, depending on the expression level of the GefA (**Fig. S2C**). In either case, a major reduction in migration speed and macropinocytosis occurred (**Fig. S2E, S2F**). Time-lapse videos of GefA-expressing cells revealed that both morphological categories of cells barely move with almost no changes in cell shape. The cells appeared “frozen” over 10 minutes, compared to the control cells which displayed frequent shape changes and movement. The differences in migration are highlighted by the color-coded overlay (**Fig. 1E**). Because of the flattened and stretched out morphology, the average height of the GefA-expressing cells was around half of the control (**Fig. 1F**), and the cell basal area and flatness increased more than two-fold (**Fig. 1G; Fig. S2A**). Quantification of cell migration speed showed that GefA-expressing cells on average migrated at 0.5-1 μm/min, while wild-type cells migrated at 6 μm/min (**Fig. 1H**). The average aspect ratio of GefA-expressing cells and wild-type cells was around 1.5, while that of the elongated fraction ranged from 2.5-4.5 (**Fig. S2B**). Because the GefA-expressing cells have two different morphologies, we suspected that it could be the heterogeneous expression differences. Consistently, as shown in **Fig. S2C**, among the GefA-expressing cells, the higher expression levels of GefA have a larger basal area and smaller aspect ratio.

To confirm that this phenotype was indeed caused by GefA rather than a secondary effect of its expression, we conducted global recruitment of GefA. We anticipated that global recruitment would not only recapitulate the overexpression phenotype but also produce a stronger effect due to its faster and more direct activation. As shown in **Fig. 1I**, global recruitment of GefA rapidly induced multiple protrusions and expanded the cell in every direction, resulting in increased basal area and immobilization (**Movie S2**), as quantified in **Fig. 1J** and **1K. T**he aspect ratio did not change significantly due to the irregular, but symmetrical, shape of GefA-expressing cells before recruitment (**Fig. S2D**). The color-coded overlay illustrates the changes in movement and cell shape before and after GefA recruitment, showing that after recruitment, the cells expanded with more protrusions in all directions, resulting in zero net movement (**Fig. 1L**). Similarly, the global recruitment of GefA expanded the Gβ-null cells in all directions, leading to an impairment in migration, and captured by color-coded images (**Fig. S1B, S1C**). Taken together, overexpressing GefA or global recruitment of GefA that increased Ras activity resulted in cell expansion and impaired cell movement, as summarized in Fig. 1M.

### Exchange Factor GefA Activates Signal Transduction and Cytoskeletal Activities

Since overexpression or recruitment of GefA led to remarkable changes in cell morphology and alterations in migration, we investigated whether the phenotypes were indeed caused by hyperactivated Ras activity. We compared Ras activity in cells with and without GefA expression using fluorescence-labelled RBD as a biosensor. As shown in **Fig. 2A**, control cells showed micron-sized, dynamic patches of RBD signals that marked protrusions, typically lasting for 1 or 2 minutes. In contrast, GefA-expressing cells displayed much larger RBD patches on the membrane that remained relatively static for about 10 minutes in this example. When normalized to the cell perimeters, the lengths of the RBD patches are increased more than two-fold in the GefA-expressing cells compared to controls (**Fig. 2B**). Occasionally, RBD waves were observed in single GefA-expressing cells, propagating across the cell and creating a broad protrusion as the wave swept to the edge (**Fig. 2A**). Such waves of Ras activity are observed in electroshock-fused giant cells but are rare in single control cells not expressing GefA. To further assess whether GefA increases events downstream of Ras activity, we used biosensor PHcrac and LimE to examine PIP3 and F-actin polymerization activity. Normally, PHcrac and LimE mirror RBD activity, occurring with protrusions, as shown in the controls (**Fig. 2C**). As expected, in GefA-expressing cells, LimE mimicked the RBD signals and indicated significantly increased actin polymerization along the cell periphery, which could account for their flattened phenotype. However, we did not observe PHcrac labeling on GefA-expressing cell membranes (**Fig. 2D**). Additionally, to confirm that GefA expression promotes actin polymerization, we fixed cells and stained them with phalloidin, an F-actin binding reagent. As shown in **Fig. 2E**, phalloidin mainly stained macropinocytotic cups in control cells, but in GefA-expressing cells, it exhibited massive staining on the entire cell membrane along with many diffuse, short filament structures in the cytosol. These results together indicate that GefA expression increases Ras activity and promotes more actin polymerization around the entire cell periphery, which likely explains the decreased migration in GefA-expressing cells, as actin polymerization everywhere prevents movement in a specific direction.

**Fig. 2.**
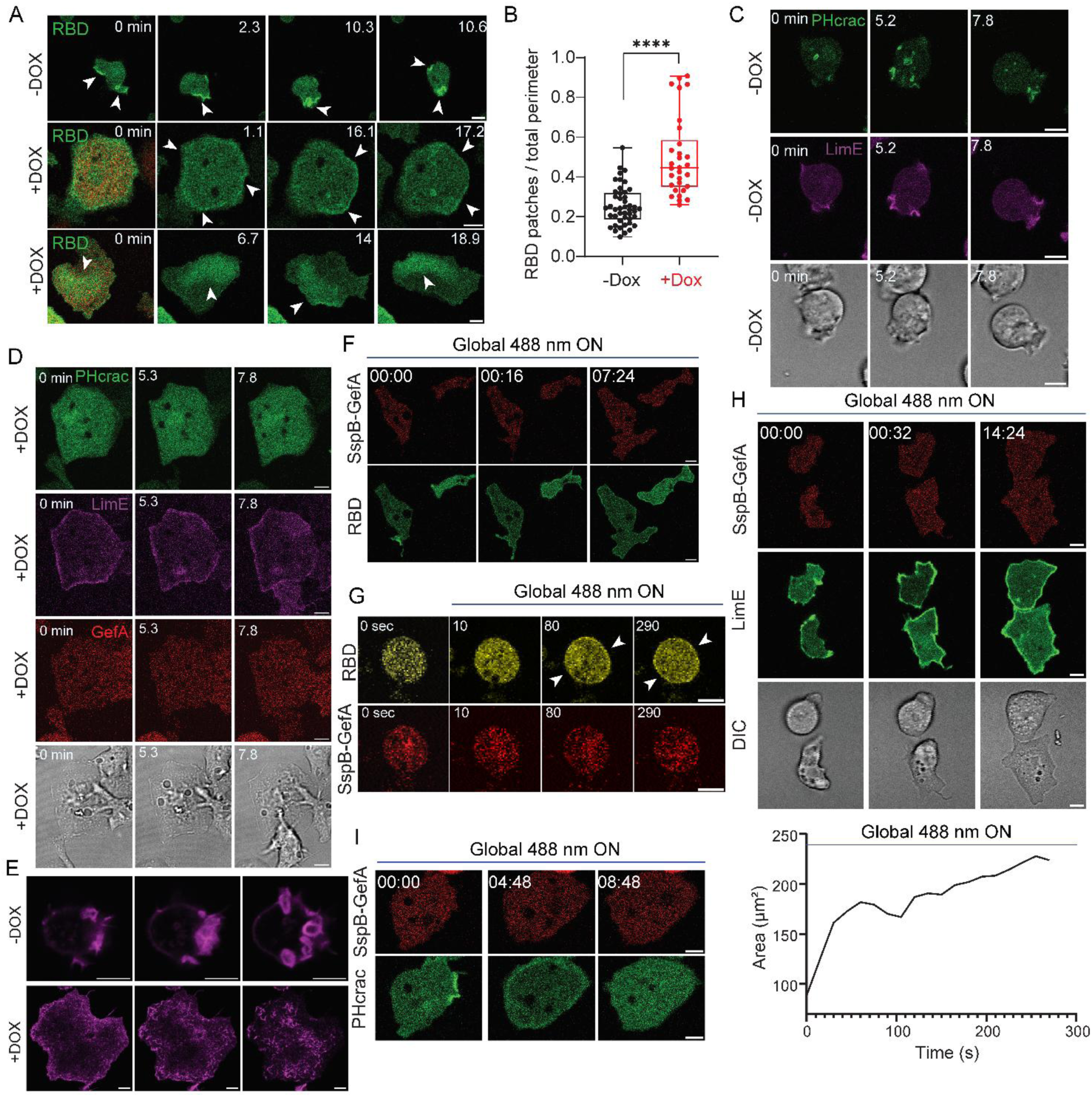
GefA activates signal transduction and cytoskeletal activities. **(A)**Confocal images of vegetative *Dictyostelium* cells expressing GFP-RBD (biosensor for activated Ras; green) minus (top) and plus (middle and bottom) induction of mRFPmars-GefA (red) expression. White arrows highlight the RBD patches (activities) in these cells. In the bottom panel, RBD shows wave-like activities. Time in minutes. Scale bars, 5 µm. **(B)** Box-and-whisker plots of the ratio between RBD patches lengths and total cell perimeter minus (black) or plus (red) GefA expression. n = 45 (−Dox) or n = 30 (+Dox) cells examined over three independent experiments; asterisks indicate significant difference, ****P ≤ 0.0001 (two-sided Mann–Whitney test). The boxes extend from 25th to 75th percentiles, median is at the center and whiskers and outliers are graphed according to Tukey’s convention (GraphPad Prism 8). Confocal images of vegetative *Dictyostelium* cells expressing PHcrac-YFP (biosensor for PIP_3_; green) and LimE-Halo (biosensor for F-actin; magenta) minus **(C)** and plus **(D)** the induction of mRFPmars-GefA (red) expression. Time in minutes. Scale bars, 5 µm. **(E)** Phalloidin staining of vegetative *Dictyostelium* cells minus (top) and plus (bottom) GefA expression, showing in three z-stacks images. Time-lapse confocal images of vegetative *Dictyostelium* cells co-expressing Opto-GefA and RBD-YFP **(F)** or LimE-YFP **(H)** or PHcrac-YFP **(I)** on 2D glass surface. Opto-GefA is recruited globally by switching on 488 nm laser. Area changes after recruitment of Opto-GefA in **(H)** is shown in line plot. Time is in min:s format. Scale bars, 5 µm. **(G)** Time-lapse confocal images of vegetative *Dictyostelium* cells co-expressing Opto-GefA and RBD-YFP in latrunculin A. The arrow shows membrane RBD patches. Time in second. Scale bars, 5 µm.

The strong actin polymerization caused by GefA overexpression was induced by overnight incubation with doxycycline, but the optogenetic recruitable GefA can promote cell expansion within minutes. We were curious to see if the fast expansion was caused by rapid signaling activity and actin polymerization downstream of Ras activation, independent of transcriptional activities. To address this question, we observed Ras activity and its downstream signaling events, PIP3 accumulation and actin polymerization directly when recruiting GefA to the cell cortex. Before recruitment, only a single RBD patch appeared at protrusions. Within 15 seconds after GefA recruitment, the entire membrane displayed RBD binding, indicating a major, global increase in Ras activity. The cell began to expand, making broader protrusions, and the RBD signals remained on the membrane throughout the recruitment period (**Fig. 2F**), even in latrunculin A treated cells (**Fig. 2G**). Concurrently, the LimE biosensor indicated that actin polymerization increased along the cell periphery, consistent with the area expansion of the cell, as quantified in the line scan (**Fig. 2H**). Actin waves were sometimes observed during this expansion, usually emerging from the center and propagating to the edge (**Movie S3**). Simultaneously, PIP3 accumulation showed a dynamic increase on the membrane. However, the PIP3 response lasted for 5 minutes before subsiding for the remainder of the time (**Fig. 2I**). The transient nature of the PIP3 response likely explains why we did not observe obvious PIP3 patches on the membrane in cells incubated overnight with doxycycline (**Fig. 2D**).

### NGAP Alters Cell Morphology and Migration

Since Ras small GTPases are regulated by both GAPs and GEFs, we examined the effects of overexpressing or recruiting RasGAPs. Previously, we found that overexpression of a C2 domain-containing RasGAP, C2GAPB, inhibited Ras activity, but when globally expressed, C2GAPB polarized cells and promoted migration(10). Building on this observation, we turned our attention to another C2 domain-containing RasGAP in *Dictyostelium*, NGAP, to investigate its role in cell migration(25).

Given the inhibitory effect of NGAP on Ras activity, we hypothesized that its local recruitment to the leading-edge protrusion would induce protrusion retraction, while its global recruitment would result either in inducing polarity as does C2GAP or whole-cell contraction and cessation of migration (**Fig. 3A**). To test this hypothesis, we illuminated a small region of the cell with blue light, thereby concentrating SspB-fused NGAP in that region. Within 30 seconds, the illuminated area of the cell rapidly contracted, causing the cell to start to migrate away from the light source (**Fig. 3B**). Next, we placed cells in microchannel devices. Consistent with the results from the 2D experiments, the enrichment of SspB-fused NGAP at the cell front induced a polarity switch (**Fig. 3C**). NGAP recruitment consistently pushed the cell to move away from the light **(Fig. 3D)**. In contrast to cells on 2D substrates, cells in confinement exhibit directed migration, which enables us to alternate their polarity repeatedly using SspB-fused NGAP (**Movie S4**).

**Fig. 3.**
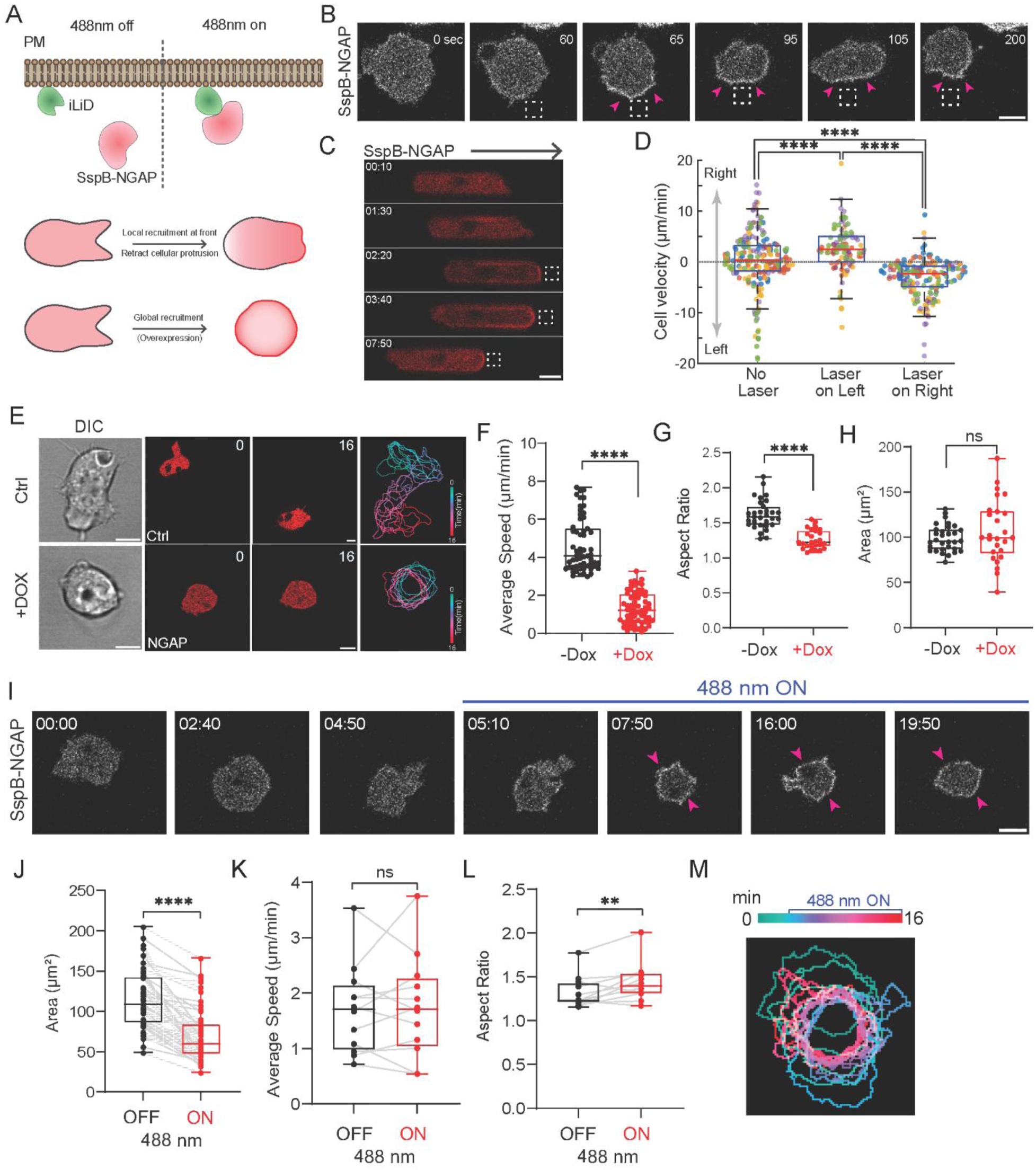
Inhibition of Ras by NGAP alters cell morphology and migration. **(A)**Schematic of the optogenetics tool. Recruitment of Opto-NGAP (mRFPmars-SspB-NGAP) to membrane-anchored iLiD upon 488 nm illumination. Typical effects at cell front or global recruitment are indicated. **(B)** Opto-NGAP recruited at one side of migrating vegetative *Dictyostelium* cell on 2D glass surface with 488 nm light (dashed white box). Pink arrows highlight the retraction of the protrusion. Scale bar, 5 µm. **(C)** Opto-NGAP recruited to the front of cell migrating to the right in a microchannel (dashed white box). Arrow indicated the migration direction before light stimulus. Scale bar, 5 µm. **(D)** Instantaneous velocity of Opto-NGAP cells when the laser light is off, to the left or right of cells. Colored dots represent each of n=6 cells; total number of measurements are 198, 152, and 152, respectively. Asterisks indicate significant difference, ****P ≤ 0.0001 (two-sided Mann–Whitney test). **(E)** Time-lapse confocal images of vegetative *Dictyostelium* cells expressing empty vector control (red; top panel) or mRFPmars-NGAP (red; bottom panel) after overnight Dox treatment. Time in minutes. Scale bars, 5 µm. DIC images (left-most panel) and overlaid outlines (color-coded 1-min intervals) of control or NGAP-expressing cells (right-most panel). Box-and-whisker plots of average cell speed **(F)**, Aspect ratio **(G)** and cell area **(H)** before (black; −DOX) or after (red; +DOX) overnight induction of NGAP. n = 31 (−Dox) or n = 27 (+Dox) cells examined over three independent experiments; asterisks indicate significant difference, ****P ≤ 0.0001; NS, not significant, P = 0.4888 (two-sided Mann–Whitney test). The boxes extend from 25th to 75th percentiles, median is at the center and whiskers and outliers are graphed according to Tukey’s convention (GraphPad Prism 8). **(I)** Snapshots of vegetative *Dictyostelium* cells expressing Opto-NGAP on 2D glass surface. Opto-NGAP is recruited globally by switching on 488 nm laser. Pink arrows highlight the contraction of the entire cell. Time in min:s format. Scale bars, 5 µm. Box- and-whisker plots of cell area **(J)**, average cell speed **(K)** and aspect ratio **(L)** before (black; 488 nm OFF) or after (red; 488 nm ON) global recruitment of NGAP. n = 50 cells (cell area) and n = 12 cells (average speed and aspect ratio) examined over three independent experiments; Statistical analysis as in **(F)** to **(H)**. NS, not significant, P = 0.9697 (two-sided Wilcoxon signed-rank test). **(M)** Color-coded (1-min interval) outlines of a representative cell before and after the presence of 488 nm light.

We further investigated the effects of overexpressing NGAP on cell morphology and migration. When NGAP expression was induced with doxycycline, cells formed tiny protrusions and rounded up, as compared to the normal-sized pseudopods in control cells with a clear front and back (**Fig. 3E**). To determine if these tiny protrusions could mediate cell migration, we recorded time-lapsed videos with and without NGAP expressions. While control cells migrated a significant distance over 16 minutes, NGAP-expressing cells hardly moved (**Fig. 3E**). The migration tracks were captured by color-coded overlay images. Quantification of average cell migration speed showed that with NGAP expression, cell migration speed was 1.5 μm/min, compared to 5 μm/min for control cells. NGAP-expressing cells also had a smaller aspect ratio but a similar basal cell area and volume when compared to control cells, indicating they were rounder and might not form large protrusions (**Fig. 3F-H**, **Fig. S4A**). Since NGAP expression reduces cell migration speed, we examined the correlation between NGAP expression levels and migration speed. As might be expected, we detected a negative correlation between NGAP intensity and cell migration speed (**Fig. S4B**), further indicating that NGAP inhibits cell migration.

Next, to directly examine the inhibitory role of NGAP on cell migration and protrusion formation, we globally recruited SspB-fused NGAP to the entire cell cortex. This recruitment of NGAP caused a dramatic whole-cell contraction within a short time (**Fig. 3I, Movie S5**). Cell area also decreased more than two-fold after NGAP recruitment (**Fig. 3J**). The cells expressing the recruitable version of NGAP already had impaired cell migration and rounded morphology before recruitment and their average migration speed and aspect ratio did not change significantly before and after recruitment (**Fig. 3K, 3L**). The whole-cell contraction after NGAP recruitment was also shown in the color-coded overlay image (**Fig. 3M)**. Based on the local, global recruitment and overexpression results, NGAP appears to inhibit cellular protrusions by inducing contraction, thereby halting cell migration. To further test this idea, we examined the effects of NGAP recruitment in myosin heavy chain null cells and RacE null cells, both of which have defects in contraction. As expected, neither local nor global recruitment of NGAP caused significant changes in area in the null mutants, compared to the two-fold changes in area reduction in wild-type (**Fig. S3A-H**; **Fig. 3I**).

### NGAP Alters Signal Transduction and Cytoskeletal Activities

Given that NGAP is a RasGAP, we examined whether the phenotypes caused by its expression or recruitment were due to decreased Ras activity. We first compared RBD signals in cells with and without NGAP expression. Control cells exhibited broad, long-lasting RBD-labeled protrusions, whereas NGAP-expressing cells only had tiny and transient RBD-labeled protrusions. The ratio of RBD patch to cell perimeter length showed that Ras activity decreases with NGAP expression (**Fig. 4A and 4B**). Similarly, we further assessed PIP3 activity and actin polymerization, which typically mirrors RBD signal patterns. As anticipated, PHcrac and LimE only transiently lit up on small puncta-like protrusions in NGAP-expressing cells, compared to multiple wide patches in control cells, clearly demonstrating that NGAP plays an inhibitory role on Ras, PI3K, and actin polymerization activities (**Fig. 4C and 4D**). Ras/PIP3 patches are also indicators of macropinocytosis; uptake measurements revealed that macropinocytosis was dramatically reduced upon NGAP expression (**Fig. S4C, S4D**).

**Fig. 4.**
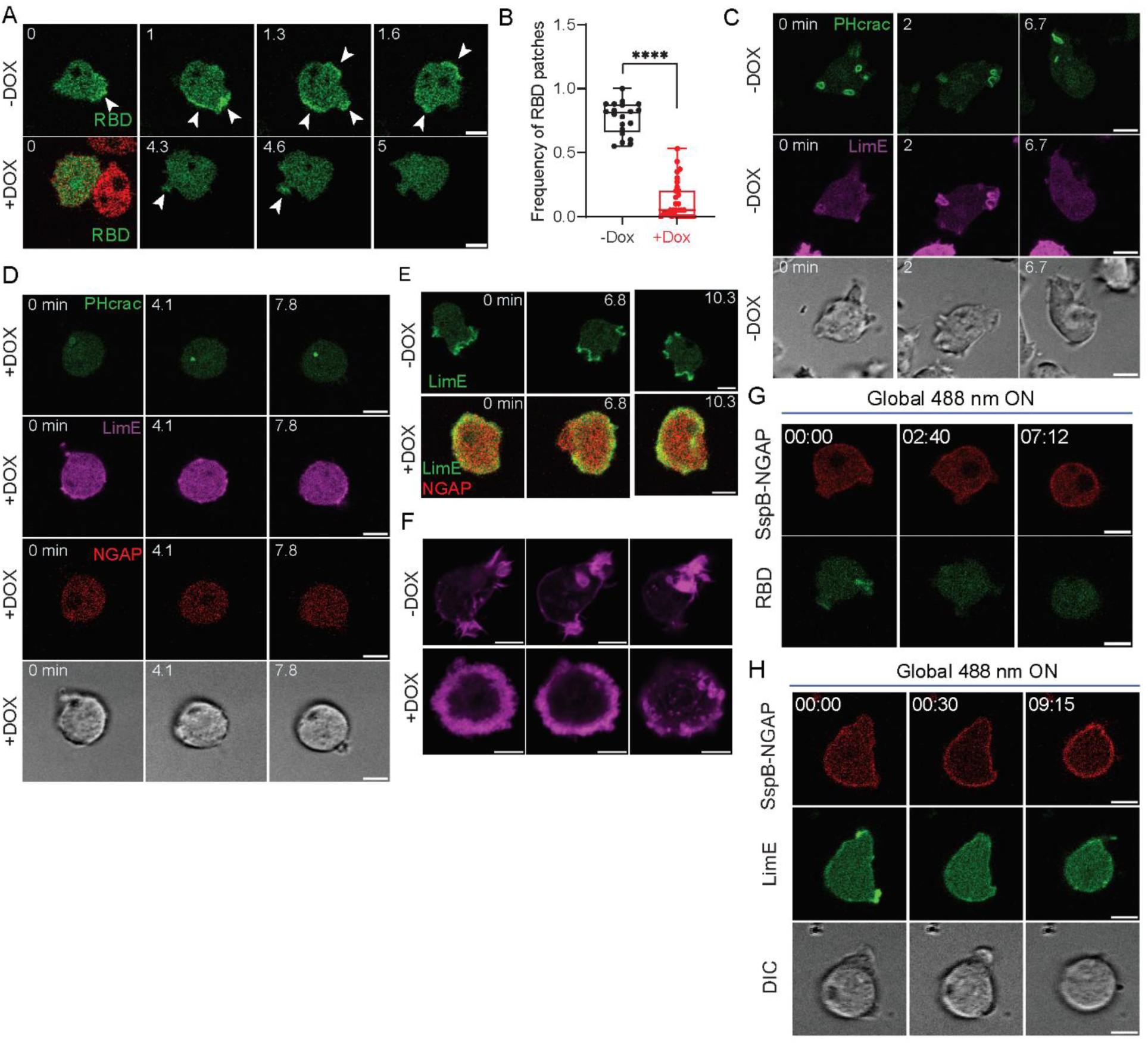
NGAP alters signal transduction and cytoskeletal activities. **(A)**Confocal images of vegetative *Dictyostelium* cells expressing GFP-RBD (biosensor for activated Ras; green) minus (top) and plus (bottom) induction of mRFPmars-NGAP (red) expression. White arrows highlight the RBD patches (activities) in these cells. **(B)** Box-and-whisker plots of the ratio between RBD patches lengths and total cell perimeter minus (black) or plus (red) NGAP expression. n = 20 (−Dox) or n = 32 (+Dox) cells examined over three independent experiments; asterisks indicate significant difference, ****P ≤ 0.0001 (two-sided Mann–Whitney test). The boxes extend from 25th to 75th percentiles, median is at the center and whiskers and outliers are graphed according to Tukey’s convention (GraphPad Prism 8). Confocal images of vegetative *Dictyostelium* single cells expressing PHcrac-YFP (biosensor for PIP3; green) and LimE-Halo (biosensor for F-actin; magenta) minus **(C)** and plus **(D)** induction of mRFPmars-NGAP (red) expression. Time in minutes. Scale bars, 5 µm. **(E)** Time-lapse confocal images of vegetative *Dictyostelium* cells expressing LimE-GFP minus (top) and plus (bottom) NGAP expression. Actin ring is visible at the bottom surface of NGAP-expressing cell. Time in minutes. Scale bars, 5 µm. **(F)** Phalloidin staining of vegetative *Dictyostelium* cells minus (top) or plus (bottom) NGAP expression, showing in three z-stacks images. Time-lapse confocal images of vegetative *Dictyostelium*cells co-expressing Opto-NGAP and RBD-YFP **(G)** or LimE-YFP **(H)** on 2D glass surface. Opto-NGAP is recruited globally by switching on 488 nm laser. Time is shown in min:s format. Scale bars, 5 µm.

Interestingly, among the NGAP-expressing population, some cells are flatter and showed LimE ring patterns around the periphery close to the bottom surface. These actin rings appeared to be “standing waves” that propagate slowly or did not propagate relative to the cell (**Fig. 4E**). This unique ring structure of actin was also confirmed by phalloidin staining in NGAP-expressing cells, which was distinct from the protrusions in control cells (**Fig. 4F**). Both small LimE-labeled protrusions and non-propagating actin rings could contribute to the decreased migration of NGAP-expressing cells.

We next tested whether NGAP functions as a RasGAP to inhibit Ras activity and its downstream PI3K and actin polymerization activity, thereby slowing down cell migration. To test this, we used our optogenetic system to globally recruit NGAP and monitored RBD and LimE signals as readouts. Before recruitment, we detected tiny RBD and LimE labeling on the small protrusions of NGAP-expressing cells. Upon blue light activation, NGAP was recruited to the cell cortex within 30 seconds, rapidly diminishing RBD and LimE signals, resulting in whole-cell contraction within 3 minutes, which was maintained for at least 10 minutes (**Fig. 4G, 4H, Movie S6**). Altogether, these results indicate that NGAP inhibits cell migration by decreasing Ras activity and actin polymerization, leading to the elimination of protrusions and cell contraction.

### Optimum Ras Activity and Cortical Tension Dictate Effective Migration

Expression of three Ras regulators, GefA, C2GAPB, and NGAP, cause either spreading, tightly localized protrusions, or tiny unfocused protrusions, which either halted migration, increased migration, or decreased migration, respectively. The shape and size of the protrusions also suggested that differences in mechanical properties, such as cortical tension, may occur.

To measure their cortical tension, we performed micropipette aspiration in GefA-, C2GAPB-, and NGAP-expressing cell lines and compared the values to those of wild type cells(34, 35). We determined the length of extension into the pipette versus the cell radius at various pressures and calculated the cortical tension (**Fig. 5B**). We observed that NGAP- and C2GAPB-expressing cells had lower cortical tension than the control cells, while GefA-expressing cells had increased cortical tension (**Fig. 5C**).

**Fig. 5.**
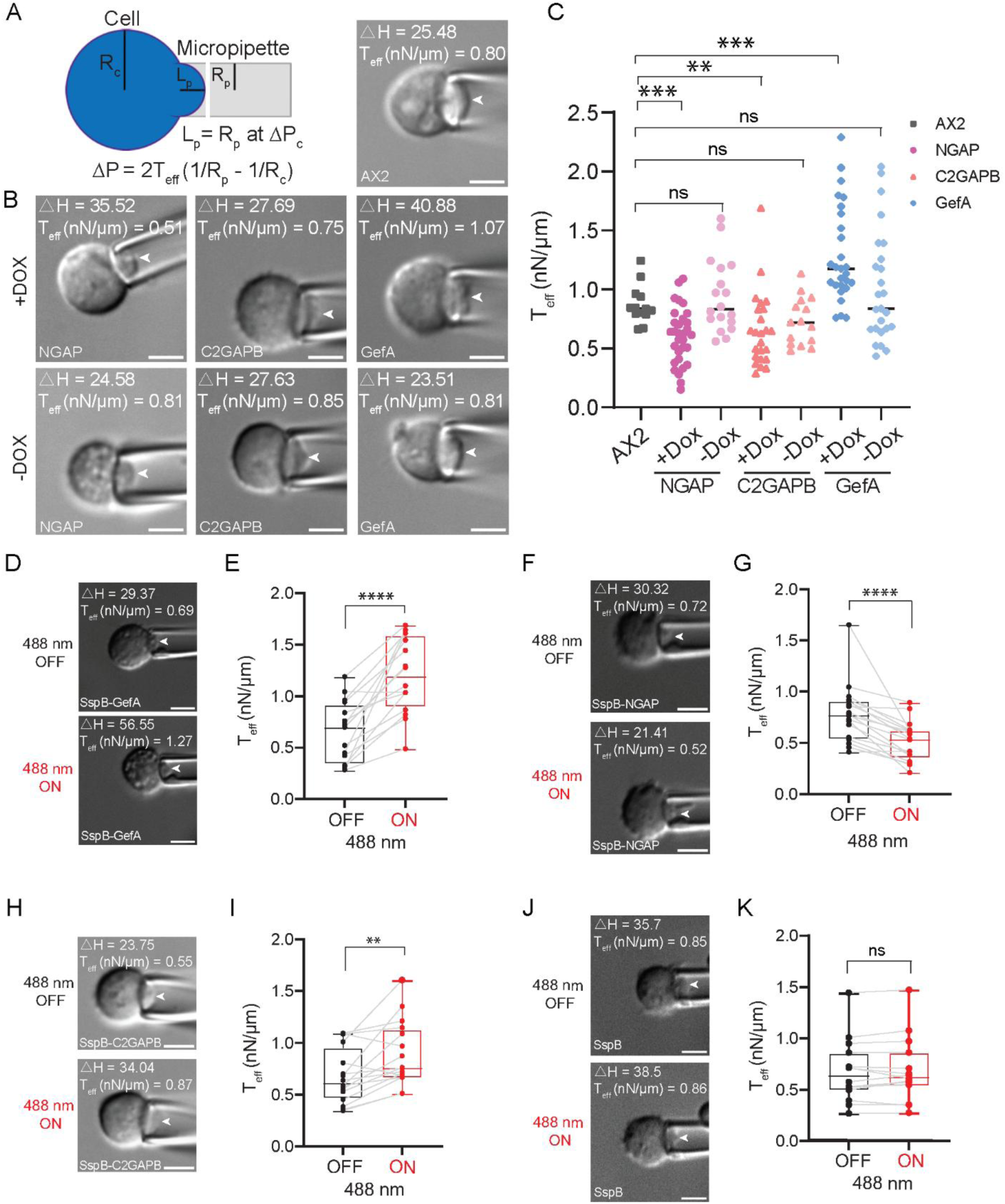
Modulation of Ras activity leads to changes in cortical tension. **(A)** Schematic of the Micropipette Aspiration (MPA) and example of MPA measurement on wild type cell (AX2, right). For cortical tension measurements, the pressure increases until the length (L_p_) of the region of the cell pulled into the micropipette equals the radius of the pipette (R_p_). At this equilibrium pressure (ΔP_c_), the radius of the cell can be measured, and the effective cortical tension T_eff_ is calculated from the equation shown in the panel. **(B)** Representative images of MPA measurements on NGAP, C2GAPB, GefA-expressing cells plus (top) and minus (bottom) the addition of DOX. **(C)** The effective cortical tension values of AX2, NGAP (± Dox), C2GAPB (± Dox) and GefA (± Dox). Measurements were quantified as detailed in the Materials and Methods. ***P ≤ 0.001; **P ≤ 0.01; NS, not significant, P = 0.9528, P = 0.1077, P = 0.7847. Cell numbers n = 13-27 from three independent experiments. P values were calculated using Kruskal Wallis (P ≤ 0.0001) followed by the two-sided Mann–Whitney test, GraphPad Prism 8. **(D)(F)(H)(J)** Representative images of MPA measurements on SspB-GefA, SspB-NGAP, SspB-C2GAPB and SspB-Ctrl cells before (top, 488nm OFF) and after (bottom, 488nm ON) recruitment. Scale bars, 5 µm. **(E)(G)(I)(K)** The effective cortical tension values of SspB-GefA, SspB-NGAP, SspB-C2GAPB and SspB-Ctrl cells before (488nm OFF) and after (488nm ON) recruitment. Measurements were quantified as detailed in the Materials and methods. ****P ≤ 0.0001; **P = 0.0027; NS, not significant, P = 0.2979. Cell numbers n = 16-18 from three independent experiments. P values were calculated using two-sided Mann–Whitney test, GraphPad Prism 8.

These initial cortical tension measurements were conducted on cell-lines that stably express the Ras regulators. In contrast, the signaling and morphological changes observed in fast moving cells are much more rapid. To test whether Ras manipulation has immediate effects on cortical tension, we leveraged our optogenetic system. Because the light-induced recruitment occurred immediately and is reversible, we decided to measure the cortical tension of the same cell twice. The first time was performed without blue light illumination. After the cell reached the critical point, we relaxed the cell and then measured the cortical tension again with blue light activation. We hypothesized that acutely altered Ras activity would augment the change of cortical tension, and as expected, recruitment of GefA further increased cortical tension by about 1.7-fold while recruitment of NGAP decreased cortical tension ∼40% from 0.8 nN/μm to 0.5 nN/μm (**Fig. 5D-G**). However, the recruitment of empty vector control had little effect on the change of cortical tension before or after recruitment (**Fig. 5J, 5K**). Because recruitment of C2GAPB increased myosin II contraction, it might lead to an increase in the cortical tension. As shown in **Fig. 5H** and **5I**, recruitment of C2GAPB slightly increased the cortical tension, but not as significantly as the changes elicited by recruiting GefA or NGAP, which could mean that C2GAPB is within the optimal range of both Ras activity and cortical tension for most effective migration.

## Discussion

In this study, we expanded our investigation into the role of Ras activity in regulating cellular migration dynamics. Using *Dictyostelium* as a model, we demonstrated that acute modulation of Ras activity via Ras GEFs and GAPs induces striking alterations in cell shape and migration behavior. By employing optogenetic tools, we replicated phenotypes previously observed in overexpression studies, but on significantly shorter timescales, highlighting the immediate influence of Ras on cell motility. Furthermore, the immediate effects of Ras modulation are independent of receptor inputs and extracellular gradients, underscoring the self-autonomously organized properties of the signaling network. Additionally, we examined how changes in Ras activity affect the mechanical properties of the cell, revealing dynamic fluctuations in cortical tension that correspond with variations in Ras signaling. Our findings establish a biphasic relationship between Ras activity levels, cortical tension, and migration efficiency, emphasizing the necessity of balanced Ras signaling and cortical tension for optimal cellular motility.

Distinct outcomes were observed between local and global recruitment of RasGEF and RasGAP. While local recruitment promoted or inhibited protrusions as anticipated, global recruitment of these Ras modulators resulted in immobilized cell movement. We propose that the amplification of Ras signaling by global recruitment pushes the system to an extreme imbalance, which results in reduction in cell motility. In our previous work, we identified an optimal level of Ras activity that maximizes cell migration (10). Building on this, our current findings further suggest a biphasic relationship between Ras activity and cell migration, where both insufficient and excessive Ras activity impair motility. Furthermore, as cortical tension has been implicated in regulating cell shape and migration behavior (36–39), we characterized the effects of our Ras modulators on cortical tension. The results showed that cortical tension parallels change in Ras activity. Together, these findings indicate the existence of a “Goldilocks”, optimal range of Ras activity and cortical tension that regulates cell behavior, as demonstrated in this study through its impact on migration.

On an unrestricted 2D glass surface, while local recruitment of RasGEF or RasGAP did induce or inhibit protrusions, cells lacked sufficient polarity to support sustained changes in cell migration. In contrast, within uncoated channels, cells migrated as polarized cells even without any external stimulus. The channel setup also allowed for more precise optogenetic manipulation of migration direction and behavior. In the channels, it was clear that local enrichment of RasGEF at the trailing edge of migrating cells in confinement was sufficient to reverse their migration directionality. Similarly, optogentic recruitment of RasGAP, albeit at the leading edge of migrating cells, changed migration direction. The effects of these distinct optogenetic stimulations on migration directionality were evident in microchannels than on 2D because of the highly directed migration induced by confining microchannel walls. Importantly, the accumulation of RasGEF or RasGAP consistently preceded the directional changes of the cells. Furthermore, even on 2D unrestricted surface, the global activation of Ras through RasGEF recruitment overrode chemotactic stimuli in wild-type cells, and local activation directly induced protrusions and polarity changes in Gβ-null cells. These findings provide strong evidence that Ras activity provides a powerful signaling input necessary for directed migration, functioning independently of receptor-mediated input.

Although Ras is generally believed to regulate cellular behavior and cytoskeletal remodeling through long-term gene regulation (40–42), our previous work and this study using optogenetics demonstrates that Ras controls cell migration over short time periods(10, 11). Notably, both cell shape and migration behaviors changed dramatically within seconds to minutes, accompanied by corresponding changes in cortical tension as Ras activity was modulated. Although cell state changes are often associated with transcriptional regulation(43), our findings highlight the effects of short-term Ras activity fluctuations that are independent of transcriptional changes. Our results indicate that these rapid, non-transcriptional mechanisms can acutely shift the cell state, potentially by driving the system away from its current setpoint. While transcriptional regulation may ultimately establish a new setpoint for long-term adaptation, this raises the question of how these immediate, non-transcriptional effects integrate and coordinate with longer-term transcriptional and translational changes to influence overall cell behavior.

Our findings have important implications for cancer treatment, suggesting that investigators targeting Ras should consider the cell’s activity state and its microenvironment. Given the biphasic relationship between Ras activity and cell migration efficiency, an ideal therapeutic approach for Ras-driven oncogenesis would aim to shift the activity level beyond the optimal range for migration, driving it toward an inactive or highly active state. (44–46). A deeper understanding of the distinct roles Ras plays in regulating cell migration, mechanical properties, and growth is crucial for the development of more targeted and effective therapeutic approaches.

## Acknowledgements

We thank all members of the Peter Devreotes, Pablo Iglesias, Douglas Robinson, and Konstantinos Konstantopoulos laboratories (Schools of Medicine and Engineering, JHU) for helpful discussions and providing resources. We thank Tian Jin (NIH), Marc Edwards (Amherst College), and Yuchuan Miao (Harvard Medical School) for providing plasmids. We appreciate DictyBase and Addgene for plasmids. This work was supported by NIH grant R35GM118177 (to P.N.D.), NIH grant S10OD016374 (to S. Kuo of the JHU Microscope Facility), NIH grants R01GM066817 (to DNR) and R01GM0149073 (to DNR and PAI), HHMI Gilliam Fellows Program GT16455 (to EP), and NIH grant R35GM156305 (to KK).

## Author Contributions

YL and PND conceived and designed the project; YL engineered constructs/stable cell lines, designed and executed experiments, and performed majority of data analyses with input from PND and PAI; YL and EP performed the MPA experiments with input from DNR; QY and SY performed the microchannel assay with input from KK; PAI and GQ assisted with some data analyses; YD performed uptake assays; JB made some constructs; PND and YL prepared initial drafts and wrote and revised final version of the manuscript with help from DNR and KK; PND supervised the study.

## Competing Interests

Robinson is co-founder of Mechanomics Discovery, LLC.

## Materials and Methods

### Reagents and Inhibitors

Hygromycin B (Thermo Fisher Scientific, 10687010) or G418 sulfate (Thermo Fisher Scientific, 10131035) was purchased as 50 mg ml^−1^ stock solution. Doxycycline hyclate (Sigma, D9891-1G) was dissolved in sterile water to make a stock of 5 mg ml^−1^. 50 mg ml^−1^ FITC-dextran (Sigma-Aldrich, 60842-46-8) was made in sterile water, and diluted to 2 mg ml^−1^ in HL5 medium. 5 mM latrunculin A (Enzo Life Sciences, BML-T119-0100) was made in DMSO. All stock solutions were aliquoted and stored at −20 °C. According to experimental requirements, further dilutions were made in development buffer (DB) before adding to the cells.

### Plasmid Construction

*Dictyostelium* NGAP (C2GAP1)-expressing plasmid was a gift from Tian Jin laboratory (NIH, Laboratory of Immunogenetics). GefA-expressing plasmid was obtained from dictyBase (cat. no. 353). Using these constructs, we subcloned NGAP or GefA into doxycycline-inducible pDM335 plasmid (dictyBase, cat. no. 523) using BglII/SpeI restriction digestion to generate mRFPmars–NGAP/pDM335, and mRFPmars– GefA/pDM335 constructs. SspB R73Q ORF was amplified from tgRFPt–SspB R73Q plasmid (Addgene #60416, RRID: Addgene_60416) and then subcloned into NGAP/pDM335 or GefA/pDM335 at the BglII site to generate the opto-NGAP (mRFPmars–SspB R73Q–NGAP) and opto-GefA (mRFPmars–SspB R73Q–GefA) constructs. tgRFPt–SspB R73Q ORF was introduced into pCV5 vector to generate the tgRFPt–SspB R73Q–Ctrl/pCV5 construct (Addgene #201761). PKBR1N150–Venus– iLiD/pDM358 construct was made previously. This plasmid was used to subclone PHcrac–YFP, LimEΔcoil–YFP or RBD–YFP ORF to generate dual expressing PKBR1N150–Venus–iLiD/PHcrac–YFP, PKBR1N150–Venus–iLiD/LimE–YFP or PKBR1N150–Venus–iLiD/RBD–YFP constructs, respectively. The shuttle vector, pDM344 (dictyBase, cat. no. 551), was used for this purpose. Constructs for GFP–RBD and GFP–LimEΔcoil were procured from the Firtel laboratory (UCSD) and Marriott laboratory (University of Wisconsin-Madison), respectively. Constructs were verified by diagnostic restriction digestion and sequenced at the JHMI Synthesis and Sequencing Facility.

### Cell Culture

WT *Dictyostelium discoideum* cells of the AX2 strain (dictyBase, cat. no. DBS0235521) was obtained from the Kay laboratory (MRC Laboratory of Molecular Biology). Gβ-null (Gβ–) cells were created in our laboratory previously. The myosin heavy chain-null strain (mhcA^−^) and RacE null (RacE^-^) were obtained from the Robinson laboratory (School of Medicine, JHU)(47, 48). All lines were cultured axenically in HL5 medium (laboratory stock) at 22 °C. Growth-stage cells were used for imaging, and developmental stage cells were used for chemotaxis assay.

### Electroporation

*Dictyostelium* stable lines were generated by electroporating 2 µg DNA into 10^7^ cells using a chilled 0.1-cm cuvette (Bio-Rad, cat. no. 1652089) at 0.85 kV/25 µF twice with a 5-s interval, followed by antibiotic selection over 3–4 weeks.

### Imaging and Microscopy

Vegetative *Dictyostelium* were adhered on an eight-well coverslip chamber for 30 min. Next, fresh DB was added to the attached cells and used for imaging. To induce GefA and NGAP expression in *Dictyostelium*, doxycycline (50 µg ml^−1^) was added the night before imaging. For macropinocytosis assay, cells were incubated with 2 mg/ml FITC-dextran for 10 min, washed twice with DB buffer and imaged subsequently. All live- or fixed-cell imaging was acquired with 0.3–0.5% laser intensity using the following microscopes: (1) a Zeiss LSM780-FCS single-point, laser scanning confocal microscope with 780-Quasar; 34-channel spectral, high-sensitivity gallium arsenide phosphide detectors supported with ZEN Black software, and (2) a Zeiss LSM800 GaAsP single-point laser scanning confocal microscope with wide-field camera supported with ZEN Blue software. All images were acquired with either 63×/1.40 PlanApo oil or 40×/1.30 PlanNeofluar oil DIC objective, along with a digital zoom.

### Optogenetics

Optogenetic experiments with vegetative *Dictyostelium* were conducted in the absence of a chemoattractant, except for chemotaxis assays. Throughout image acquisition, a solid-state laser (561 nm excitation and 579–632 nm emission) was used for visualizing proteins or recruitable effectors fused to an mRFPmars or tgRFPt tag. Images were acquired for 5–10 min, after which a 450/488 nm excitation laser was switched on globally to activate recruitment. Image acquisition and photoactivation were performed at ∼15-s intervals. Using the T-PMT associated with the red channel, we acquired DIC images. The interactive photobleaching module on a Zeiss LSM800 was used to perform local recruitment experiments. A small region of interest (ROI) was placed near migrating cells, and bleached with a 488 nm laser (laser power of ∼0.5% for *Dictyostelium*) in single iteration. The time interval of photoactivation and image acquisition was ∼15 s.

### Microfluidics device fabrication, cell seeding and optogenetic control

Microfluidic devices composed of polydimethylsiloxane (PDMS) with an array of parallel microchannels (200 µm in length, 3 μm in width, and 3 μm or 10 μm in height) were fabricated as previously described(49). A laser profilometer was used to verify the height and width of the microchannels. GefA-expressing cells were tested in narrow (3 × 3 µm²) channels, while NGAP-expressing cells were examined in 3 × 10 µm² channels.

Cells were detached from culture plates, followed by centrifugation at 300 g for 5 min and resuspension in DB buffer. 20 μL of cell suspension was added into the cell-seeding inlet to establish a pressure-driven flow that directed cells into the microfluidic channels. Cells were allowed to adhere and spread at the microchannel entrances for 5 min before all inlets and outlets were filled up to 100 μL with DB buffer. Then time-lapse images were recorded in a Nikon AXR confocal microscope using a Plan Apo 40×/1.15 WI (AXR) objective.

Cells migrating within confining channels were monitored in real-time by imagining with 561nm laser to identify their leading and trailing edges. Optogenetic stimulation was administered using a 488 nm laser at 1% power for 1 second within a defined rectangular region, targeting either the leading or trailing edge of the cell. Stimulations were repeated at 10 sec intervals for 5 min to enable consistent localization of GefA or NGAP on the membrane.

### Micropipette aspiration (MPA) and effective cortical tension quantification

The MPA setup was described in the previous review(35). Micropipettes of ∼5 μm diameter (exact diameter was measured for each pipette during use) were stabilized at the bottom of the cell chamber. Exponential growth phase cells were seeded in imaging chambers at ∼10% confluency. Small aspiration pressure was generated to form attachment between the cell and the pipette tip. The cell was then raised off the surface, and the aspiration pressure was increased to the equilibrium pressure (ΔP) at which the length of the cell inside the pipette (L_p_) is equal to the radius of the pipette (R_p_). Effective cortical tension was quantified using the Young-Laplace equation:

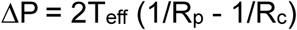

where ΔP = aspiration pressure that produced a deformation where L_p_ = R_p_, T_eff_ = effective cortical tension, R_p_ = radius of the pipette, and R_c_ = radius of the cell outside the pipette.

### Image Analysis

All images were analysed with Fiji/ImageJ 1.52i (NIH), MATLAB (Mathworks), and Python v.3.10. We utilized GraphPad Prism 8 (GraphPad), MATLAB (Mathworks), and Microsoft Excel (Microsoft) for plotting our results.

To segment the cells, images were first binarized using ImageJ, by adjusting threshold to cover all pixels of the wave or cell. The range was not reset and the ‘Calculate threshold for each image’ option was unchecked. Subsequently, using the ‘Analyze Particle’ function, a size-based thresholding was applied (to exclude non-cell particles) and cell masks were generated. Next, ‘Fill holes’, ‘Erode’ and ‘Dilate’ options were applied, sequentially and judiciously, to obtain the proper binarized mask for cells. The segmented cells were used for cell outline overlays. For RBD patches quantification, the length of RBD patches and the total cell perimeter were segmented with Fiji line tool and calculated the ratio. Cell speed, area and aspect ratio quantifications were described previously(10). Macropinocytosis uptake was analyzed by outlining each cell and quantifying the total FITC fluorescence signal within each outline divided by the cell area. Data for Fig. 1D and 3D were obtained by custom code in MATLAB. Cells were segmented using thresholding, followed by filling holes (imfill) and morphological closing (imclose). The ‘regionprops’ command was used to find the cell centroid. The centroid of the laser was obtained similarly. Instantaneous speeds were obtained by one step differences and separated depending on whether a laser was present or whether it was to the left or right of the cell. Statistical test was done using a Mann-Whitney test in MATLAB.

### Statistics and Reproducibility

If datasets were normally distributed, statistical analyses were executed using unpaired or paired two-tailed nonparametric tests on GraphPad Prism 8. Results are expressed as mean ± s.d. from at least three independent experiments. NS, P > 0.05, *P ≤ 0.05, **P ≤ 0.01, ***P ≤ 0.001, ***P ≤ 0.0001. Tukey’s convention was used to plot box-and-whisker plots. For non-normally distributed datasets, for groups >2, the Kruskal Wallis test was used followed by pairwise Mann Whitney. Mann Whitney alone was used if datasets only had two groups. Statistical test details are indicated in the figure legends. No statistical methods were used to predetermine sample sizes, but our sample sizes are similar to those reported in previous publications.

### Manuscript Writing

ChatGPT v.4o software (https://chat.openai.com/chat) and Gemini 2.0 flash (https://gemini.google.com/app) were employed in the writing of the Results and Significance section. Each figure was manually examined and a series of bullet points explaining each panel of the figure was written by the authors. Then ChatGPT and Gemini were asked to convert the bullet points to better sentences with correct grammar. Next, the text was carefully reviewed by each author where incorrect sentences were removed. Finally, the manuscript was run on several plagiarism software platforms which returned no instances.

---

**Fig. S1.**
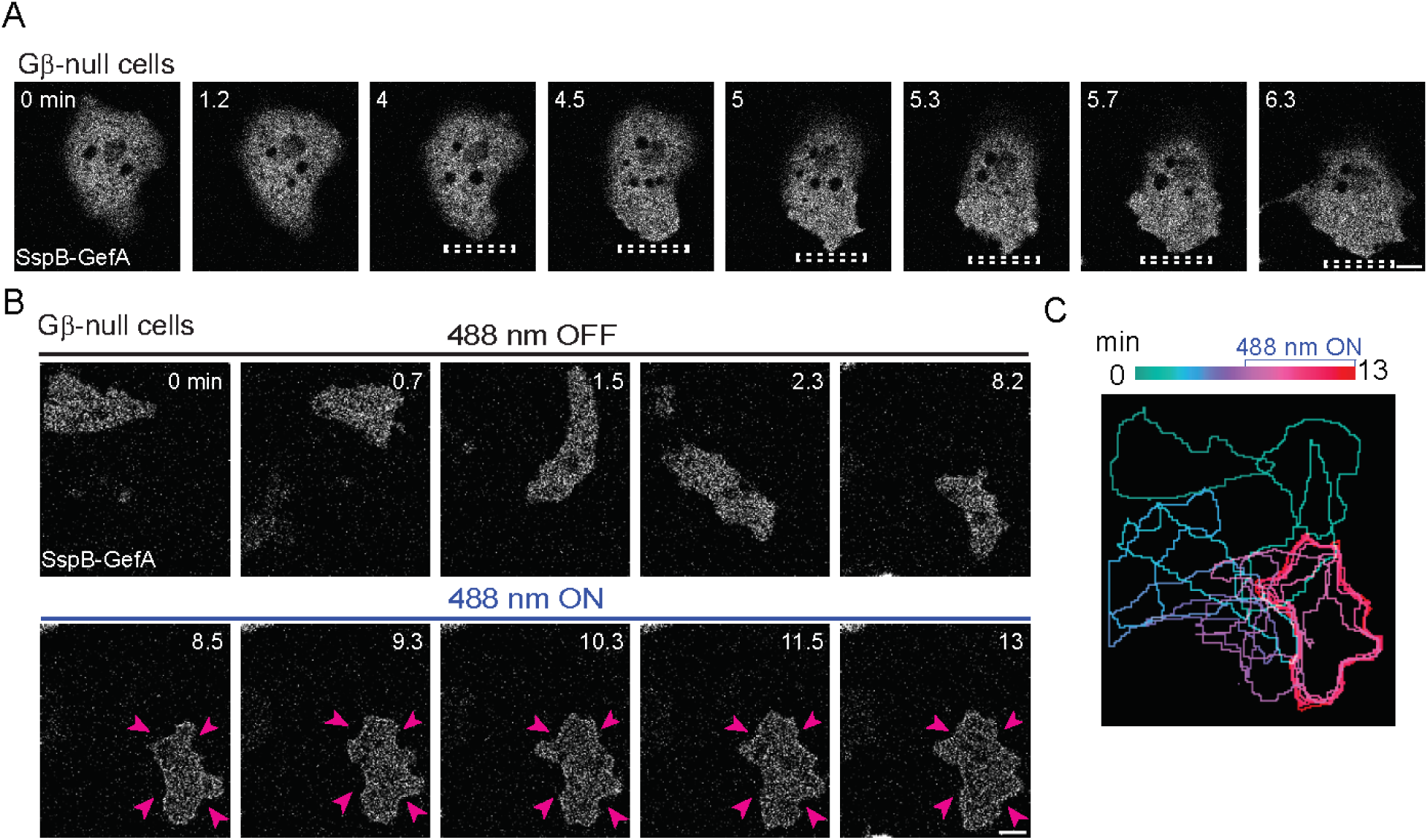
Recruitment of GefA alters Gβ-null cell morphology and migration. **(A)** Opto-GefA recruited near one side of migrating vegetative Gβ-null *Dictyostelium* cells on 2D glass surface with 488 nm light (dashed white box).Time in minutes. Scale bars, 5 µm. **(B)** Opto-GefA recruited globally in vegetative Gβ-null *Dictyostelium* cells on 2D glass surface by switching on 488 nm laser. Pink arrows highlight the expansion of the entire cell. Time is shown in minutes. Scale bars, 5 µm. **(C)** Color-coded (1-min interval) outlines of the representative cell in **(B)** before and after the presence of 488 nm light.

**Fig. S2.**
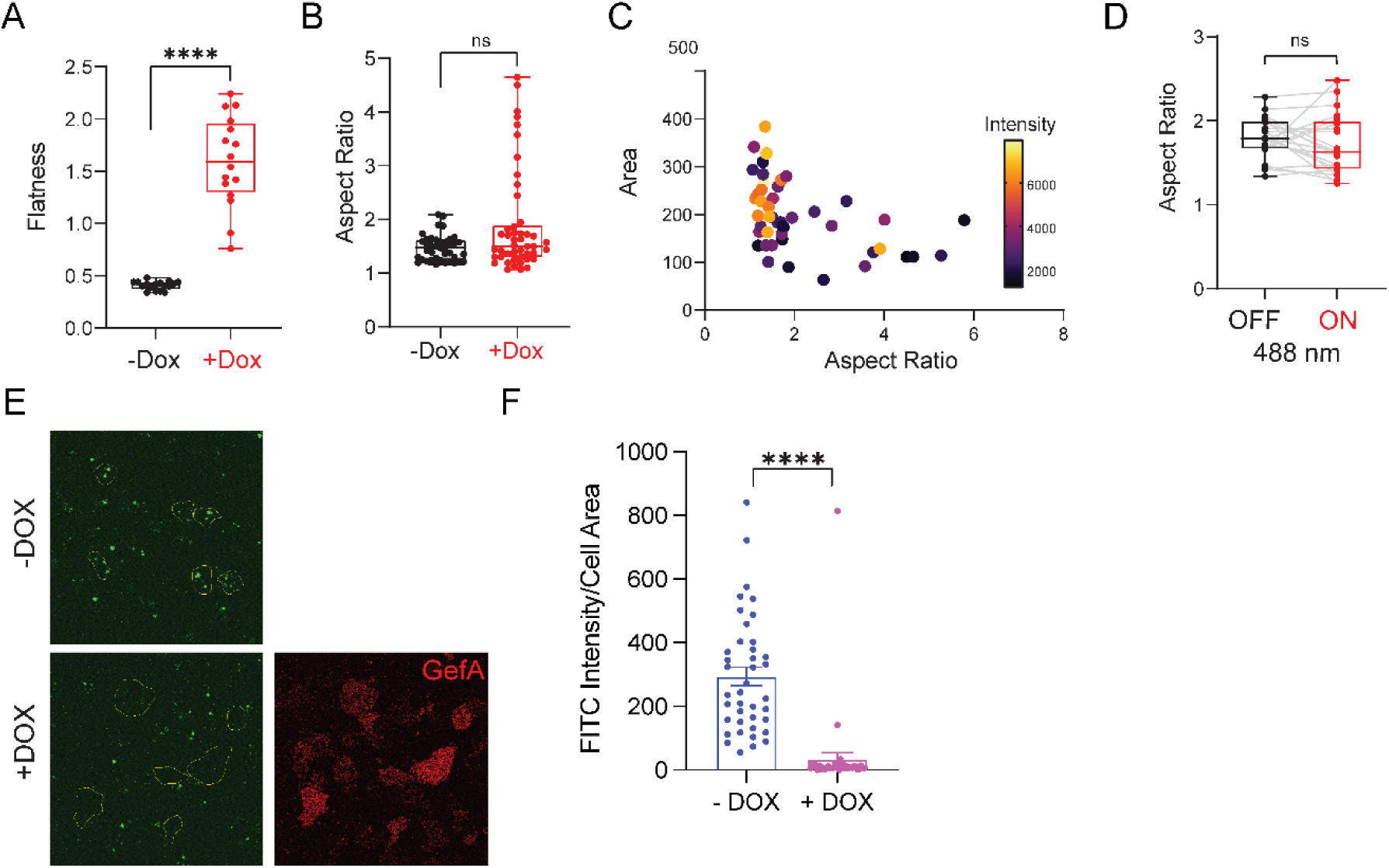
The effects of GefA on cell volume, aspect ratio and macropinocytosis. **(A)**Box-and-whisker plots of Flatness minus (black; -DOX) or plus (red; +DOX) overnight induction of GefA. n = 17 (−Dox) or n = 16 (+Dox) cells examined over three independent experiments; asterisks indicate significant difference, ****P ≤ 0.0001 (two-sided Mann–Whitney test). **(B)** Box-and-whisker plots of Aspect ratio minus (black; - DOX) or plus (red; +DOX) overnight induction of GefA. n = 40 (−Dox) or n = 49 (+Dox) cells examined over three independent experiments; asterisks indicate significant difference, NS, not significant, P = 0.877 (two-sided Mann–Whitney test). **(C)** Scatter plot of Area VS Aspect ratio with Expression Intensity color-coded scatter. n = 47 cells examined over three independent experiments. **(D)** Box-and-whisker plots of aspect ratio before (black; 488 nm OFF) or after (red; 488 nm ON) global recruitment of GefA. n = 21 cells examined over three independent experiments; NS, not significant, P = 0.2157 (two-sided Wilcoxon signed-rank test). **(E)** Representative images of vegetative *Dictyostelium* minus (top panel, -DOX) and plus (bottom panel, +DOX) doxycycline-induced mRFPmars-GefA (red) expression treated with FITC-dextran (green) before imaging. Scale bars, 10 µm. **(F)** Macropinocytosis uptake measurements, minus (blue) and plus (pink) GefA expression. n = 39 cells for both ‘-DOX’ and ‘+DOX’, over 3 independent experiments; asterisks indicate significant difference, ****P ≤ 0.0001 (Two-sided Mann-Whitney test). For all Box-and-whisker plot, the boxes extend from 25th to 75th percentiles, median is at the center and whiskers and outliers are graphed according to Tukey’s convention (GraphPad Prism 8).

**Fig. S3.**
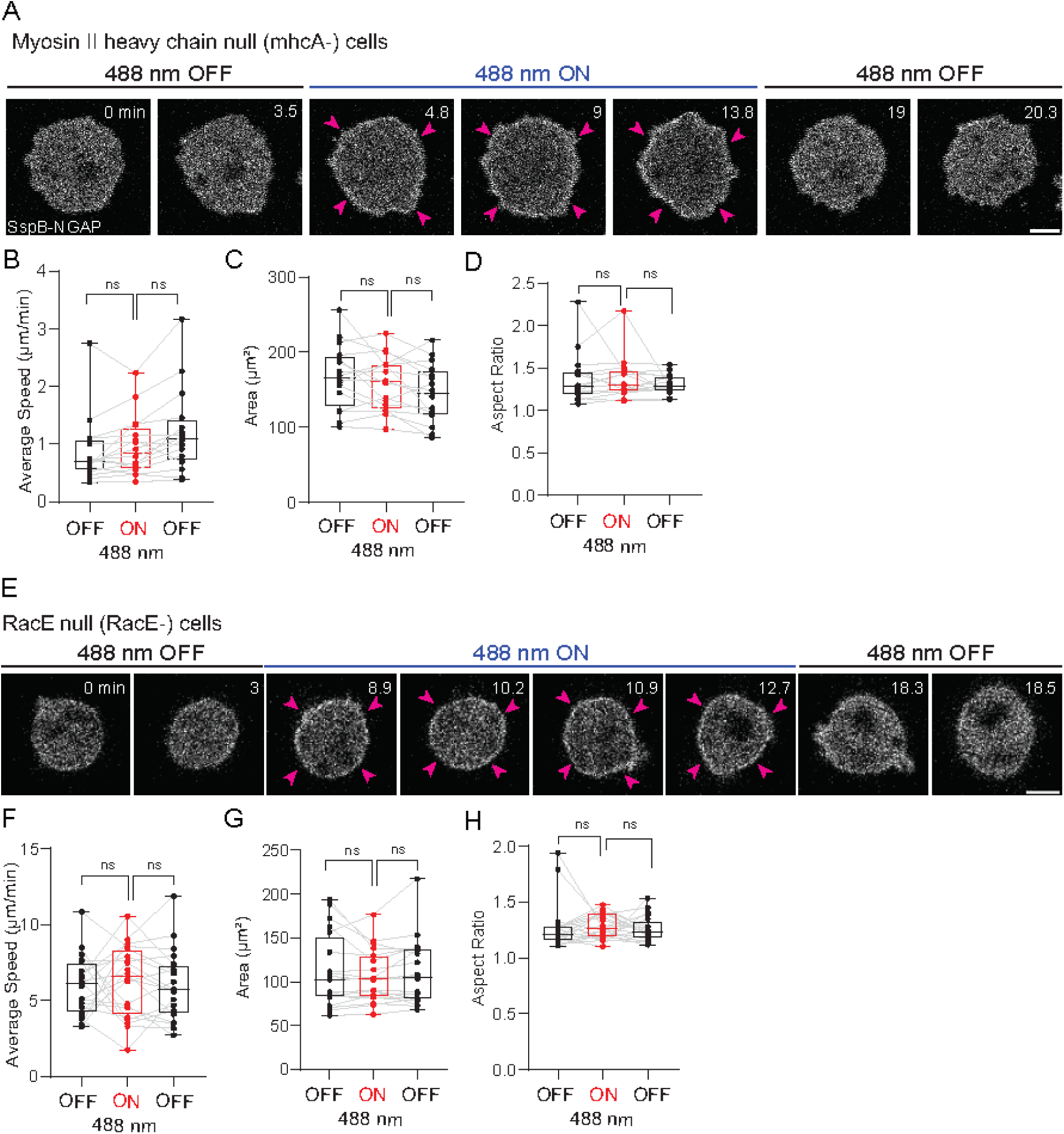
Recruitment of NGAP does not alter mhcA^-^ and RacE^-^ cell morphology and migration. Time-lapse confocal images of vegetative Myosin II heavy chain null (mhcA^-^) **(A)** and RacE null (RacE^-^) **(E)** cells *Dictyostelium* cells expressing Opto-NGAP on 2D glass surface. Opto-NGAP is recruited globally by switching on 488 nm laser and allowed to return to cytosol by switching off 488nm laser. Pink arrows highlight the membrane recruitment of Opto-NGAP but no significant changes on cell shapes. Box-and-whisker plots of average cell speed **(B, F)**, cell area **(C, G)**, and aspect ratio **(D, H)** before (black; 488 nm OFF) or during (red; 488 nm ON) and after (black; 488 nm OFF) global recruitment of NGAP. n = 16 cells (mhcA^-^) and n = 20 cells (RacE^-^) examined over three independent experiments; asterisks indicate significant difference, **P= 0.0076 **(B)**, *P= 0.0335 **(C)**; NS, not significant, P = 0.1167 **(B)**, 0.2979 **(C)**, 0.2744 and 0.8603 **(D)**, 0.7012 and 0.4524 **(F)**, 0.7841 and 0.2774 **(G)**, 0.2774 and 0.2943 **(H)** (two-sided Wilcoxon signed-rank test). The boxes extend from 25th to 75th percentiles, median is at the center and whiskers and outliers are graphed according to Tukey’s convention (GraphPad Prism 8).

**Fig. S4.**
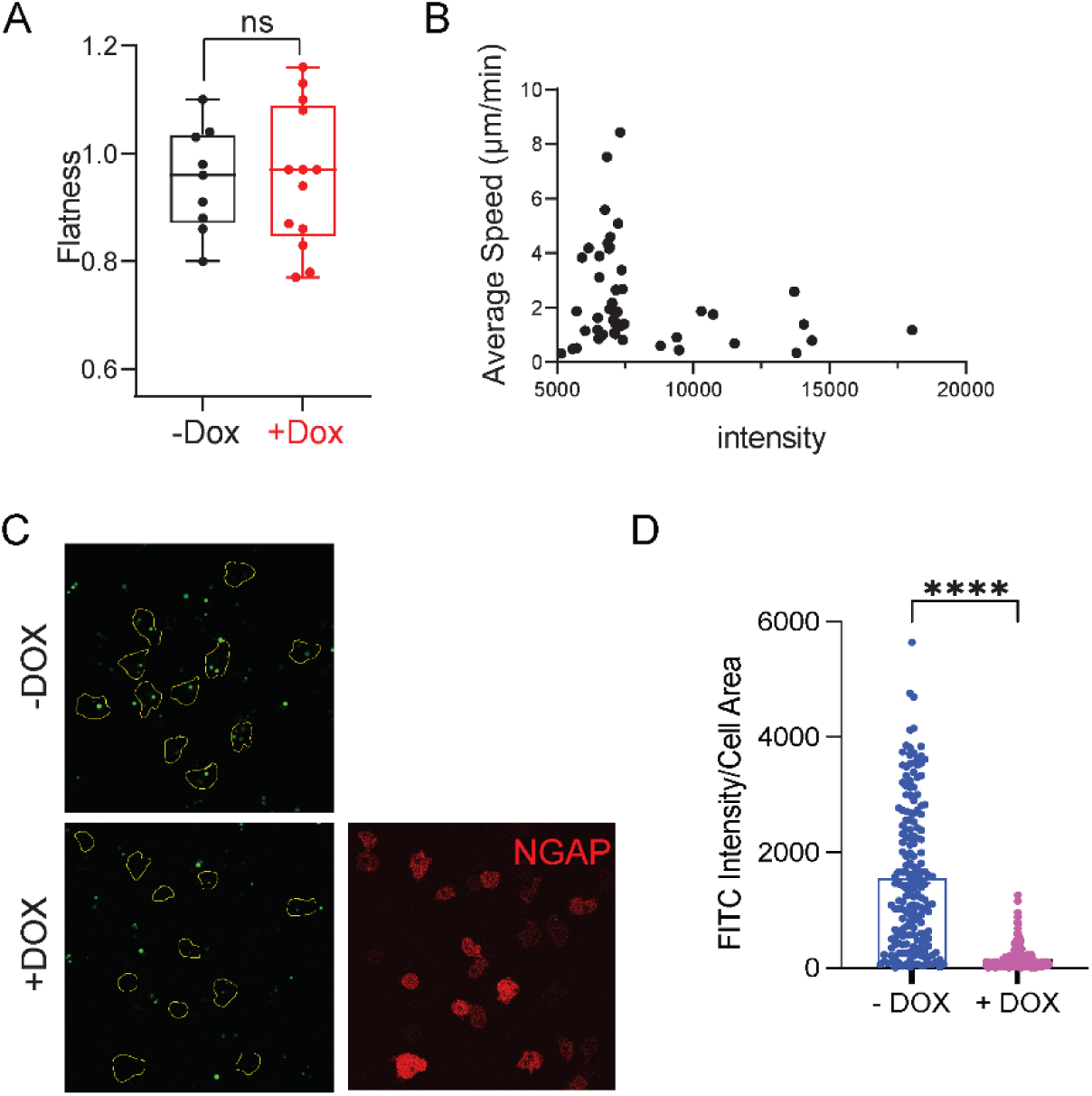
The effects of NGAP on cell volume, aspect ratio and macropinocytosis. **(A)**Box-and-whisker plots of Flatness minus (black; −DOX) or plus (red; +DOX) overnight induction of NGAP. n = 9 (−Dox) or n = 13 (+Dox) cells examined over three independent experiments. NS, not significant, P = 0.9871. (two-sided Mann–Whitney test). **(B)** Scatter plot of Average Speed VS Expression Intensity. n = 58 cells examined over three independent experiments. **(C)** Representative images of vegetative *Dictyostelium* minus (top panel, -DOX) and plus (bottom panel, +DOX) doxycycline-induced mRFPmars-NGAP (red) expression treated with FITC-dextran (green) before imaging. Scale bars, 10 µm. **(D)** Macropinocytosis uptake measurements, minus (blue) and plus (pink) the induction of NGAP expression. n = 182 cells for ‘-DOX’ and 203 cells for ‘+DOX’, over 3 independent experiments; asterisks indicate significant difference, ****P ≤ 0.0001 (Two-sided Mann-Whitney test). Boxes extend from 25th to 75th percentiles, median is at the center, and whiskers and outliers are graphed according to Tukey’s convention (GraphPad Prism 8).

Movie S1

Local recruitment of Opto-GefA (red) by red box to a migrating cell in a microchannel. Top left corner shows time in min:s format. Scale bar, 5 µm.

Movie S2

Global recruitment of Opto-GefA (red) to a migrating cell on 2D surface. Global recruitment was initiated when blue laser was applied globally (‘488 nm on’ at top of the video) at ‘00:50’. Top left corner shows time in min:s format. Scale bar, 10 µm.

Movie S3

Time-lapse confocal images of vegetative *Dictyostelium* cells co-expressing Opto-GefA (red; left) and LimE-YFP (green; middle) on 2D glass surface. Global recruitment was initiated when blue laser was applied globally (‘488 nm on’ at top of the video) at ‘00:00’. Top left corner shows time in min:s format. Scale bar, 10 µm.

Movie S4

Local recruitment of Opto-NGAP (red) by red box to a migrating cell in a microchannel. Top left corner shows time in min:s format. Scale bar, 5 µm.

Movie S5

Global recruitment of Opto-NGAP (grey) to a migrating cell on 2D surface. Global recruitment was initiated when blue laser was applied globally (‘488 nm on’ at top left of the video) at ‘05:00’. Top left corner shows time in min:s format. Scale bar, 5 µm.

Movie S6

Time-lapse confocal images of vegetative *Dictyostelium* cells co-expressing mRFPmars-NGAP (red; middle) and LimE-GFP (green; left) on 2D glass surface. Top left corner shows time in min:s format. Scale bar, 5 µm.

